# Ready-to-use nanopore platform for ethanolamine quantification using an aptamer-based strand displacement assay

**DOI:** 10.1101/2023.02.27.530168

**Authors:** Isabel Quint, Jonathan Simantzik, Lars Kaiser, Stefan Laufer, Rene’ Csuk, David Smith, Matthias Kohl, Hans-Peter Deigner

**Affiliations:** Institute of Precision Medicine, Furtwangen University, Jakob-Kienzle-Strasse 17, Villingen-Schwenningen, 78054, Germany; Institute of Pharmaceutical Sciences, Department of Pharmacy and Biochemistry, Eberhard-Karls-University Tuebingen, Auf der Morgenstelle 8, Tuebingen, 72076, Germany; Tuebingen Center for Academic Drug Discovery & Development (TüCAD2), 72076 Tuebingen, Germany; Institute of Organic Chemistry, Martin-Luther University Halle-Wittenberg, Kurt-Mothes-Str. 2, 06120 Halle (Saale), Germany; Fraunhofer Institute IZI (Leipzig), Perlickstrasse 1, 04103 Leipzig, Germany; Fraunhofer Institute IZI (Leipzig), Schillingallee 68, 18057 Rostock, Germany; Faculty of Science, Eberhard-Karls-University Tuebingen, Auf der Morgenstelle 8, Tuebingen, 72076, Germany

**Author notes:** both authors contributed equally (Isabel Quint, Jonathan Simantzik).

## Abstract

In recent decades, nanopores have become a promising diagnostic tool. Protein and solid-state nanopores are increasingly used for both RNA/DNA sequencing and small molecule detection. The latter is of great importance because small molecules are difficult or expensive to detect using available methods such as HPLC or LC-MS. Moreover, DNA aptamers are an excellent detection element for sensitive and specific detection of small molecules. Here, we describe a method for the quantification of ethanolamine using Oxford Nanopore’s ready-to-use sequencing platform. To this end, we have developed a strand displacement assay using a binding ethanolamine aptamer and magnetic beads. The displaced aptamer can be detected using the MinION® nanopores and analysed/quantified using our in-house developed analysis software.

## Background

The detection of small molecules is of great importance for many applications in molecular diagnostics, drug development or disease monitoring, but it is a major challenge. Their small size makes specific antibody binding difficult because only one epitope is present [5]. State-of-the-art methods include HPLC, LC/MS, or GC/MS [9, 30, 53], but these are expensive and require trained personnel for their analysis [56]. Nanopores as analytical sensors for small molecules have become increasingly recently. By measuring the current interruptions that occur during the molecule translocation through the nanopore, the analyte can be detected [42]. Nanopores are classified into three types: (1) biological nanopores embedded in a lipid bilayer; (2) synthetic nanopores fabricated in solid substrates such as Si3N4, Al2O3, TiO2, and graphene; and (3) hybrid nanopores, protein pores embedded in synthetic membranes [19]. Modified, protein or solid-state nanopores have been used several times to detect small molecules [4, 17, 43, 58]. However, the fabrication of solid-state nanopores with constant diameters is difficult and costly. On the other hand, the stability of protein nanopores is often insufficient. In addition, many pores are specifically modified for the detection of a particular molecule and are therefore unsuitable for the detection of other analytes. Moreover, these specially designed nanopores can often be used only once or are difficult to reconstruct [2, 54].

The MinION® is a pocket-sized device developed in 2014 as the first commercially available sequencer that uses nanopore sequencing technology for nucleic acid analysis [52]. Its commercially available flow cells contain an artificial membrane with up to 2048 embedded modified protein pores arranged in 126-512 controllable channels. The sensor detects minute changes in current as nucleic acids pass through the pores, this results in different signals depending on the sequence. These “events” are recognized by the integrated software (MinKNOW®) as a sequence of 3-6 nucleotide long k-mers [24]. To date, there are only two published articles from the same research group using the MinION® to detect small molecules via unassisted voltage-driven translocation [25, 50]. For this, they used complementary oligos labeled with osmium to detect their target DNA or RNA sequence (e.g., microRNA) [25]. However, to date, there is no label-free approach for short DNA or RNA sequences.

A closely related problem is the availability of easy-to-use solutions for data analysis. Since the instrument was designed specifically for sequencing, most of the tools available for handling MinION® data are geared towards sequencing analysis only. The usual type of data generated also does not include the raw current traces but consists only of reads that have already been filtered and pre-processed by the instrument, which does not provide meaningful results in the case of non-sequencing research questions. However, ONT offers an experimental mode for the MinION® that allows to acquire the complete raw signals in a so-called bulk file format. For the analysis of such files, there are some dedicated software solutions such as BulkVis [40] or SquiggleKit [12], but they only aim at sequencing analysis and are therefore not applicable to our data besides just visualizing the signal. Kang et al. provide osbp_detect [14], a software that accompanies their analysis, but its main purpose is to count the occurrence of the events they defined, which did not prove helpful with our data.

Aptamers represent a promising alternative to antibodies due to their stability, rapid binding kinetics and reversibility of binding. Aptamers are short single-stranded DNA or RNA sequences, usually <100 nucleotides (nt), that can specifically bind small molecules. Most aptamers originate from the SELEX process. The concept of the SELEX process is based on the ability of these small oligonucleotides to fold into unique 3D structures that can interact with a given target molecule with high specificity and affinity [47]. Aptamers have already been selected against a variety of targets, such as metal ions [22, 41], antibiotics [55], proteins or peptides [11, 13, 57], and other small biomarkers [7, 21, 45]. To demonstrate the principle, we chose the small metabolite ethanolamine, for which there is a well-described binding aptamer [21, 34]. Ethanolamine is an important analyte for environmental chemistry and life sciences. As a substance widely distributed in nature, it plays an important role in the healthy metabolism of humans and plants [29, 36, 39] Due to its ability to sequester CO_2_ from volatile gases, it can also contribute to climate change mitigation [8]. In the human body, ethanolamine is mainly present in the form of phosphatidylethanolamine (PE). The average level of free ethanolamine in blood is 2 µM (range 0-12 µM) in a healthy adult, while it is higher in other body fluids such as breast milk (46.2 +/- 18.1 µM), cerebrospinal fluid (14.1 +/- 3.0 µM), or saliva (135.99 +/- 96.22 µM). Newborns also have significantly higher blood levels of 52.3 µM (range 26.2-91.7 µM) [23]. Decreased PE and thus ethanolamine levels have been shown to contribute to the development and severity of various diseases such as Alzheimer’s disease [16], Parkinson’s disease [35], Huntington’s disease [10] and mitochondrial dysfunction [37]. Therefore, ethanolamine may play an important role as a potential biomarker for early disease detection. In this work, we demonstrate the ability of the MinION® sequencer to detect and further quantify ssDNA aptamers with the goal of developing a label-free system for indirect quantification of associated small targets using ethanolamine. In addition, we have developed our own visualization and analysis software called Nanotrace [15], which allows us to view actual traces of fast5 files and perform signal analysis and quantification of our aptamer data.

## Results and Discussion

### Aptamer detection with MinION®

As shown in Figure 1A, the ethanolamine aptamer exhibited a significant current interruption from 240 pA to 60, 80, or 100 pA during molecular translocation. The density plot (Figure 1B) shows the characteristic wave-like pattern of the ethanolamine aptamer. Compared to ONT flush buffer as a negative control, no significant peaks and only a small amount of noise can be seen here (see Figure 1B left). The density plot shows the distribution of the signal in the respective pico-ampere regions over the entire period of the run. The EA aptamer shows three peaks in the density plot corresponding to the pore stages: 1. base current/open pore (200-250 pA) 2. event current/aptamer translocation (60-100 pA) and 3. closed pore (0 pA). For 1 µM EA aptamer, the mean signal density was 0.322 +/- 0.117 SD. The ethanolamine aptamer contains 42 nucleotides forming a quadruplex tertiary structure [21]. This structure is bulkier and therefore leads to a longer translocation time of the ethanolamine aptamer compared to linear DNA strands of the same nucleotide size. This could be an explanation for the relatively long translocation time of the aptamer of 1 to 2 seconds, while according to literature ssDNA or RNA strands with less than 100 nucleotides translocate too fast (5 to 10 µs per base) to be detected by the MinION® with a sampling rate of 3.012 kHz. Therefore, most other methods for detecting e.g. microRNA involve deep bioinformatics or base modification to increase the signal [26, 50, 51]. The wavy pattern of our signal in the density plot results from the three different current interruptions at 60, 80, or 100 mV during aptamer translocation. This suggests that our aptamer is either in different confirmations or leads to different blocking of the nanopore depending on which side of the bulky complex passes through the pore first [1, 3, 28, 32]. Nevertheless, Figure 1 clearly shows that the ethanolamine aptamer can be detected using the MinION® nanopores.

**Figure 1:**
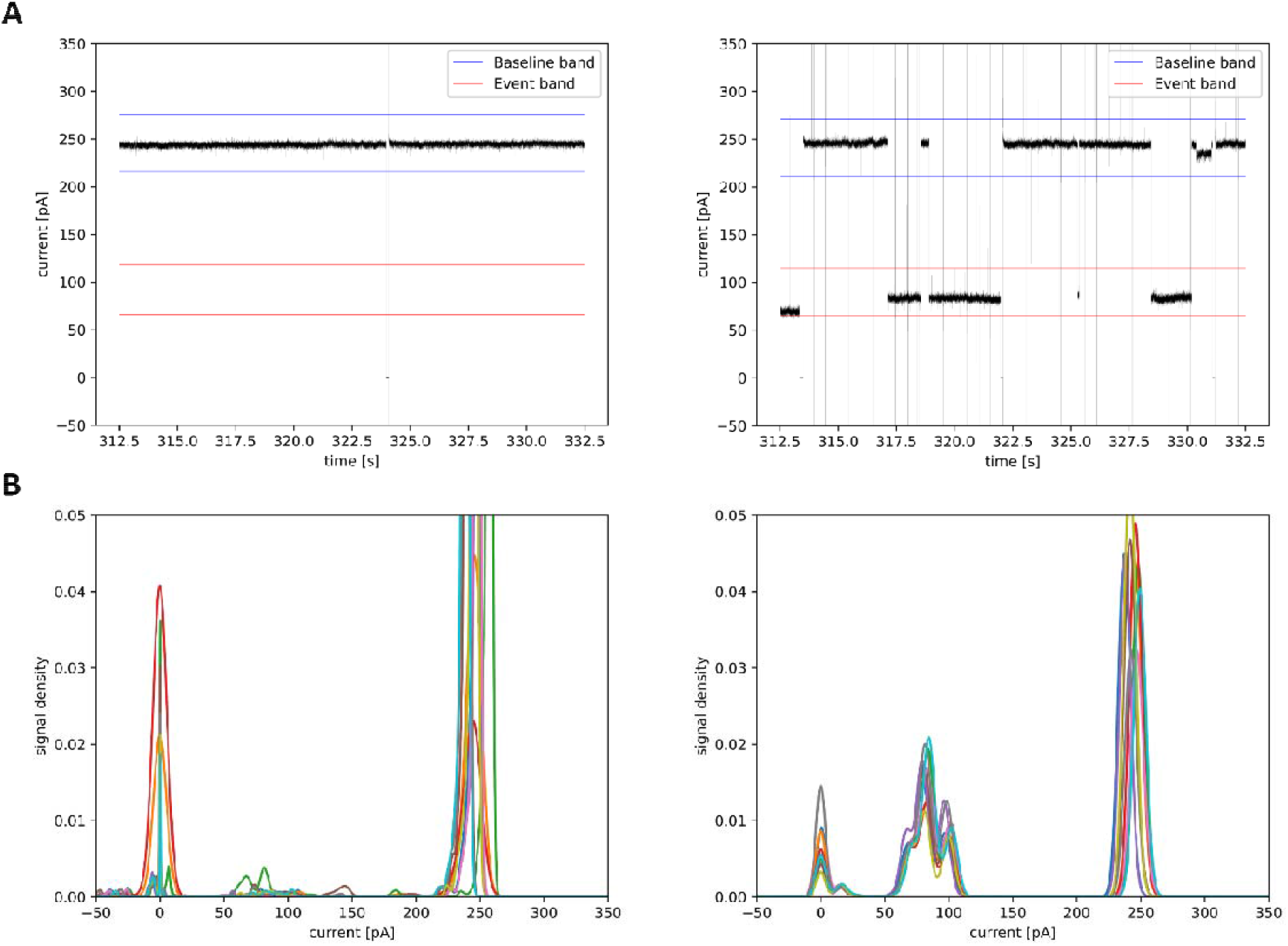
(**A**) current traces of MinION®-nanopores with ONT flush buffer (left) and 1 µM ethanolamine aptamer (right). (**B**) density plots of 1 µM ethanolamine aptamer (right) vs. ONT-flush buffer (left). The density plot shows the distribution of the signal in the respective pA ranges over the entire period of the run.

Furthermore, the ethanolamine aptamer can not only be detected but also quantified with the MinION®. As shown in Figure 2A, higher aptamer concentrations lead to an increased signal density, which is concentration dependent up to 2.5 µM. A further increase in aptamer concentration did not result in a further increase in signal density. Therefore, the linear range of aptamer detection is between 0 and 2.5 µM. This saturation effect could be explained by the availability of free pores. If all pores are permanently blocked by translocating aptamer, no higher translocation rate per time unit can be achieved. Here, the maximum translocation rate was between 30 and 40 aptamers per minute. At this translocation rate, the pores are blocked with aptamers 70% of the time (see Figure 2A). In addition, the limit of detection (LOD) was set at 50 nM because lower concentrations were not significantly different from the negative control. This is a signal-to-noise ratio issue due to the instrument and the buffer used. If the signal-to-noise ratio could be increased, the LOD could be decreased. We tried different buffer systems, but the ONT-flush buffer still showed the best signal-to-noise ratio. By calculating the standard deviations (SD), we created a calibration line with triplet measurements from each aptamer concentration (see Figure 2B). The standard deviations, especially for the lower aptamer concentrations, were quite high and also contributed to increased LOD. Nevertheless, this calibration curve is suitable for quantifying unknown aptamer concentrations and therefore served as the basis for our strand displacement assay. Furthermore, the results show that label-free detection and quantification of small oligonucleotides with nanopores is possible, which has not been described before.

**Figure 2:**
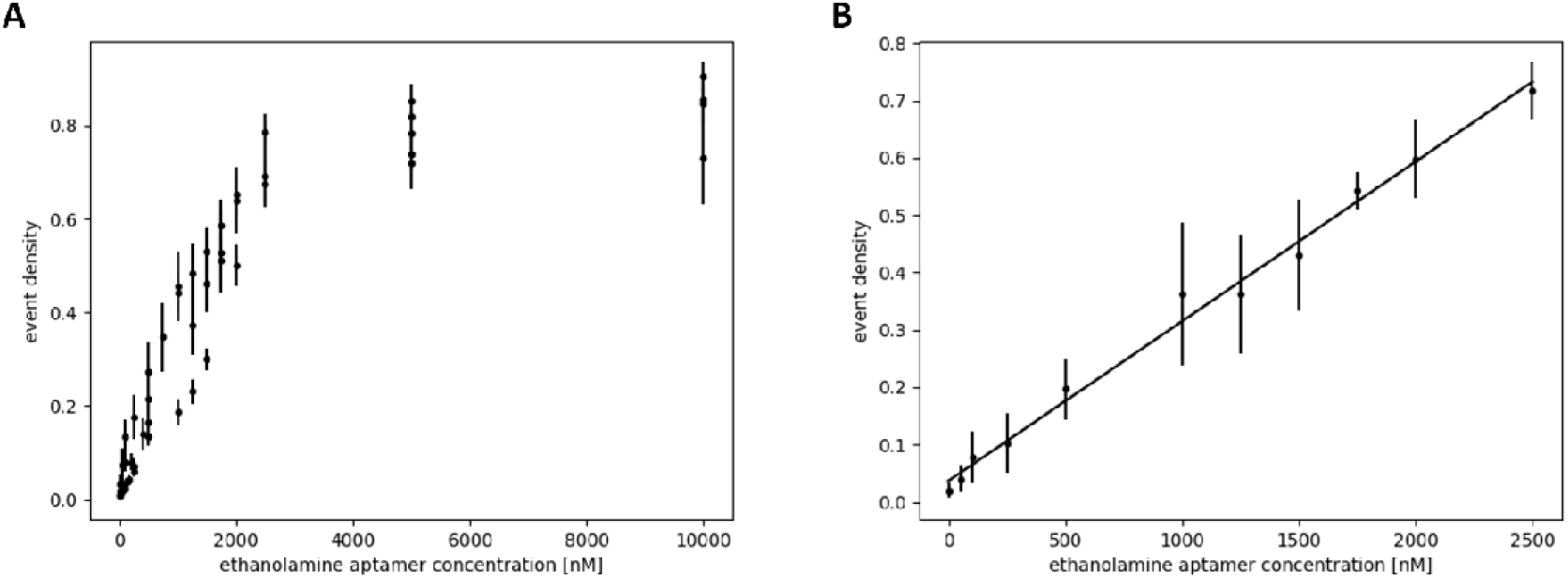
(**A**) graph of mean event densities for different aptamer concentrations. Here each replicate is plotted separate, the error bars depict the standard deviation between the active channels. (**B**) calibration line of the linear range between 0 and 2.5 µM for EA-aptamer. Therefore, the mean event densities and SD of n=3 replicates were calculated.

### Strand-displacement assay

Figure 3 shows the principle of our strand displacement assay, in which we indirectly measure the aptamer displaced by ethanolamine. This technique was chosen because, as described in the previous section, the flow cells of the MinION® showed the best signal-to-noise ratio when using the ONT flush buffer. However, this buffer has an inappropriate salt concentration and pH for aptamer complement strand binding [45]. Hybridization experiments in flush buffer showed very low binding affinity of aptamer to its complementary strand. Moreover, we observed an almost complete release of the hybridized aptamer from its complement when the buffer was changed from W&B to flush buffer after hybridization. Therefore, we decided to redesign the assay and measure the remaining bead-bound aptamer rather than the displaced aptamer (see Figure 3). The displaced aptamer, and thus the ethanolamine concentration, can then be calculated by subtracting the amount of remaining bead-bound aptamer from the negative control without ethanolamine. Both aptamer binding and ethanolamine displacement were performed using the manufacturer’s recommended W&B buffer. In this buffer, the aptamer had a high binding efficiency to its complementary strand. The binding efficiency was measured using implen’s nanodrop photometer. The best binding efficiency was obtained at a 2:1 ratio of aptamer to its complementary strand, which resulted in a mean aptamer binding of 273 pmol of aptamer per 500 µl of beads (1mg/ml). This is consistent with the manufacturer’s claim that 1 mg of beads can bind up to 500 pmol of biotinylated DNA oligos. Thus, in the W&B, the aptamer binding efficiency was 100% or even slightly higher. Higher binding efficiencies may result from non-specific binding of the aptamer to the bead surface or from slight measurement inaccuracies of the instrument since the aptamer is very short and low concentrated at 42 nucleotides. Non-specifically bound aptamer could be removed by subsequent washing.

**Figure 3:**
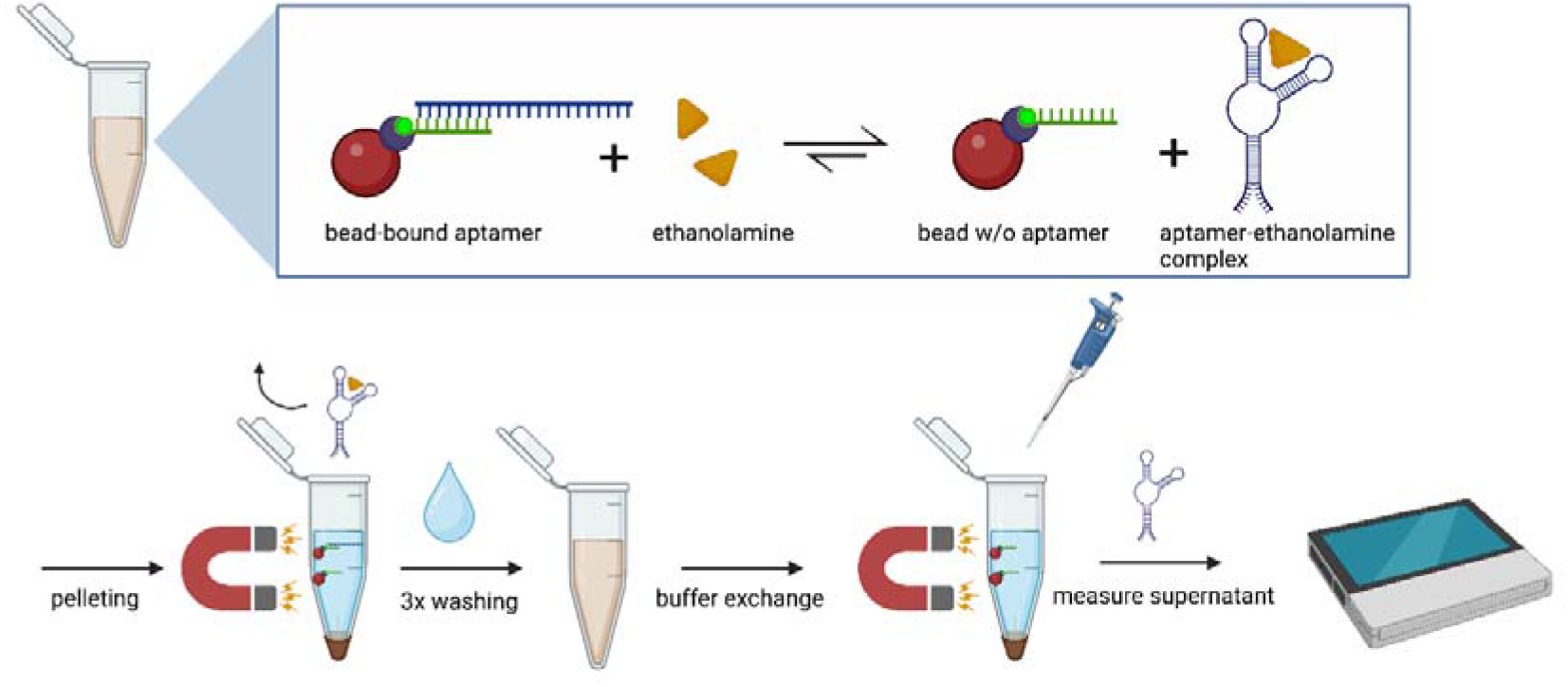
Principle of the strand displacement assay. The biotinylated ethanolamine aptamer binding complement is covalently attached to streptavidin coupled magnetic beads and the ethanolamine aptamer is hybridized to its complementary strand. When ethanolamine is added, the aptamer is displaced from its complementary strand to form the ethanolamine-aptamer complex. Beads can be pelleted with a magnet and the supernatant containing the displaced aptamer can be removed. After washing, the remaining bead-bound aptamer can be removed from the beads with ONT-Flush buffer and the amount of aptamer in the supernatant can be measured using the MinION®-device.

In Figure 4 shows the results of the strand displacement assay with different ethanolamine concentrations. As the ethanolamine concentration increases, the remaining aptamer concentration on the beads decreases as more aptamer is detached from its complement to form the aptamer-target complex with ethanolamine (Figure 4A). Saturation was reached at 20 µM ethanolamine, as higher concentrations did not result in further aptamer displacement. The maximum amount of displaced aptamer (1 µM) was reached with 20 µM ethanolamine, which means that a 20-fold excess of ethanolamine is required to displace all aptamer from the beads (see Figure 4B). However, even lower concentrations show a steadily increasing amount of displaced aptamer that can be measured. The LOD of the system was determined at 5 µM ethanolamine, as the SDs of lower concentrations are too high to be significantly different from the negative control. Aptamer concentrations were calculated from the calibration straight line (see Figure 2B) of our previous experiments by substituting the measured event densities into the straight-line equation (see Figure 4B). The displaced aptamer was calculated by subtracting the amount of aptamer bound to the beads after ethanolamine addition from the amount of aptamer bound to the beads before ethanolamine addition (Figure 4C). These results demonstrate that the MinION® instrument is suitable for indirect measurement of ethanolamine concentration. Although other methods showed lower LODs for ethanolamine detection [21, 31, 33], our assay system is sufficiently sensitive for most clinical applications. Serum concentrations of ethanolamine are around 2 µM (range 0-12 µM) in healthy patients, while they are higher in other body fluids such as breast milk (46.2 +/- 18.1 µM), cerebrospinal fluid (14.1 +/- 3.0 µM), or saliva (135.99 +/- 96.22 µM). Neonates also have significantly higher blood levels of 52.3 µM (range 26.2-91.7 µM) [23]. Ethanolamine contributes to the formation of phosphatidylethanolamine (PE), a membrane phosphatide that plays an important role in almost all cell membranes. Decreased levels are associated with several diseases such as Alzheimer’s disease, Parkinson’s disease, and Huntington’s disease. Serum levels of PE and free ethanolamine were significantly decreased in diagnosed Alzheimer’s disease patients at all stages of dementia, and disease severity correlated with dementia severity. Similar results were found in patients with Parkinson’s disease and Huntington’s disease [10, 16, 35]. Other studies showed that reduced cerebrospinal ethanolamine levels (<12.1 µM) correlate with major depressive disorder (MDD) and could be used as a biomarker for a subtype of MDD [38]. In the gut, ethanolamine plays a role in the bacterial population as some pathogenic bacteria (e.g., EHEC) can utilize ethanolamine as a carbon/nitrogen source, resulting in a competitive advantage over other bacteria in the microbiome. For example, it has been shown that virulence gene expression in EHEC can be activated with as little as 1 µM ethanolamine [27]. In another metabolomic study, mass spectrometry was used to determine that the concentration of ethanolamine in the saliva of patients with pancreatic cancer was significantly increased compared to healthy patients [49]. Increased PE concentrations were also observed in pancreatic and breast cancer cells [48]. Therefore, the sensitivity of our assay in the millimolar range is useful for monitoring ethanolamine levels in various diseases and provides an opportunity to further investigate the function of ethanolamine in these diseases without the need for complex and expensive equipment. On the other hand, our assay is just one example of how the MinION® can be used to detect small molecules with aptamers. The methodology can be transferred to other small molecules with a suitable aptamer of similar shape/size. However, we have clearly demonstrated that the ethanolamine aptamer can be quantified using the MinION® nanopores without prior labelling. Previous approaches did not manage without labelling. The long translocation time of 1-2 seconds observed here is likely due to the tertiary structure of the aptamer. Since this structure is also formed by other aptamers, our approach offers the possibility to detect other small molecules based on their aptamers [44]. However, further studies are needed for this purpose.

**Figure 4:**
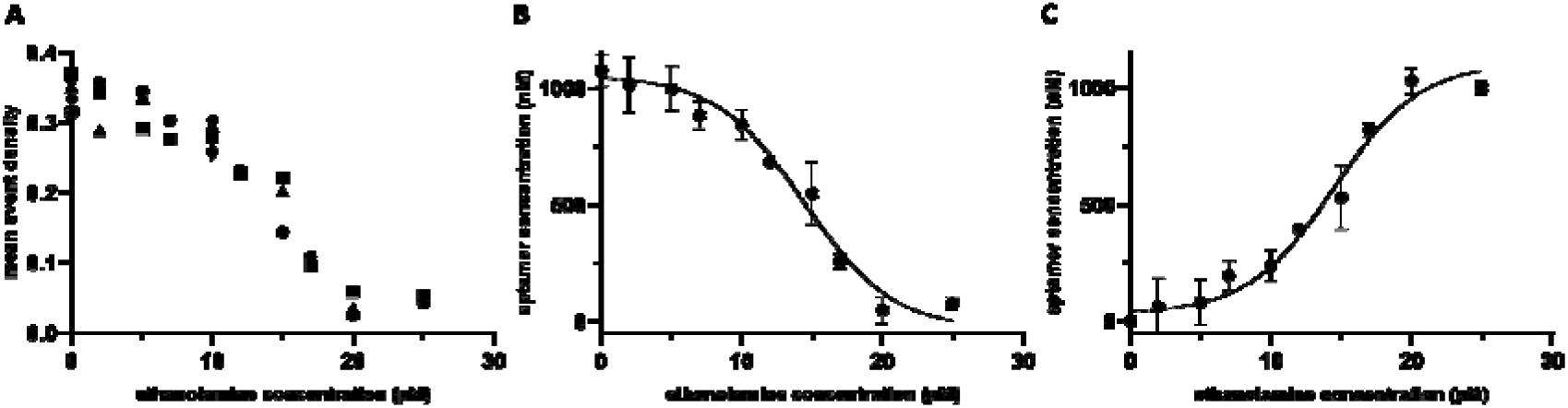
Results for the ethanolamine aptamer displacement experiment. **(A)** calculated mean event densities of non-displaced aptamers at increasing ethanolamine concentrations using MinION®. Mean event densities were calculated using Nanotrace. **(B)** Conversion of mean event densities to non-displaced aptamer concentrations (µM). Therefore, the mean signal density values from Figure 5 A were substituted into the aptamer calibration line equation (see Figure 2 B). **(C)** Displaced aptamer (µM) with increasing ethanolamine concentrations (SD and non-linear regression). The displaced aptamer was calculated by subtracting the amount of aptamer bound to the beads after ethanolamine addition from the amount of aptamer bound to the beads before ethanolamine addition.

### Data analysis

The Nanotrace software displays the squiggle plot of the selected channel and can calculate the corresponding signal density, where the concrete event boundaries can be freely selected to allow easier application in new contexts. In addition, an overlay of the event densities of 10 channels can be displayed graphically. The software has a user-friendly interface and can be used without programming knowledge. Therefore, our software allows scientists without a bioinformatics background to easily and quickly visualize the generated fast5 files. Compared to other available software tools, this is a great advantage, as most tools for evaluating bulk files use a Python-based interface and require extensive programming skills [12, 40]. Detailed information can be found in the appendix.

## Conclusion

Our results show that the MinION® device is not only suitable for DNA/RNA sequencing, but that its nanopores can also be used for the detection and quantification of small molecules. This involves the use of current-independent, controlled translocation of target-binding aptamers. In our work, we have shown an example of how such detection of small molecules might work by performing a strand displacement assay for ethanolamine with its binding aptamer. Our results demonstrate that ethanolamine can be indirectly detected by measuring its binding aptamer using MinION® nanopores with a sensitivity in the micromolar range. This is a promising approach for future clinical applications, as the MinION® device is a cost-effective and already established ready-to-use platform. The detection of small molecules, including ethanolamine, is still difficult and expensive so far, but plays an important role in the diagnosis of many diseases, as ethanolamine is associated with Alzheimer’s disease, Parkinson’s disease or Huntington’s disease. Moreover, we have developed a user-friendly open-source evaluation software (Nanotrace) to detect and quantify our ethanolamine aptamer using the fast5 files generated by MinKnow®. This method is only a proof of principle that can be adapted to other small molecules with a binding aptamer of similar structure and represents a milestone for future experiments.

## Materials and methods

### Materials

Ethanolamine binding aptamer (5’ ATACCAGCTTATTCAATTTGAGGCGGGTGGGTGGGTTGAATA 3’) and 3’-biotinylated aptamer complement (5 ‘CCACCCACCC 3’/biotin) were synthesized and purified by Integrated DNA Technologies (Coralville, IA, http://www.idtdna.com). Ethanolamine was purchased from Sigma-Aldrich (Darmstadt, Germany, www.sigmaaldrich.com). Magnetic particles 1µM (Dynabeads™ MyOne™ Streptavidin C1 product number: 65002) were purchased from Thermo Fischer Scientific (Sankt Leon-Roth, Germany, www.thermofischer.com). Flongle® flow cells and ONT-flush buffer were purchased from oxford nanopore technologies (Oxford, UK, www.nanoporetech.com).

## Methods

### Strand displacement Assay

Streptavidin-coupled Dynabeads® 1 μm magnetic beads were diluted to 1 mg/ml^−1^ in 1x washing and binding buffer (W&B-buffer: 5 mM Tris-HCl, 0.5 mM EDTA, 1M NaCl, pH 7.5) and used at this concentration throughout all experiments. The coupling procedure was performed according to the manufacture protocol as follows: a volume of 50 μL (10 mg/ml^−1^) particles was diluted in 450 μL W&B-buffer in an 0.5 ml Eppendorf tube. The particles were washed a total of 3 washes with W&B-buffer. Afterwards, a volume of 5 μL of the buffer was replaced by 5 μL 3’-biotin labelled aptamer complement at a concentration of 100 μM and incubated 90 minutes at room temperature on a shaker. The particles were washed again 3 times with W&B-buffer, to remove unbound aptamer complement. The ethanolamine aptamer was hybridized to its complement, by replacing 10 µl buffer with 10 µl 100 µM ethanolamine aptamer solution. The hybridization was allowed to proceed on a shaker at room temperature for another 90 minutes. The particles were washed again 3 times with W&B-buffer and the first supernatant was collected for evaluating binding efficiency. For strand-displacement, 0-25 µM ethanolamine in W&B-buffer was added to the beads and incubated 15 min at room temperature on a shaker. The particles were washed again 3 times with W&B-buffer and directly resuspended in 100 µl of ONT-flush buffer. After another 15 min of incubation on a shaker, 30 µl supernatant were measured on the MinION®.

### Quantification of oligonucleotides immobilization and hybridization efficiency

Nanophotometer (Nanophotometer® P-class, P-360, Implen GmbH) was used to determine the amount of immobilized aptamer; 1 μl of the supernatant was transferred to the nanodrop and a triplicate measurement were recorded per sample. The same measurements were done with non-reacted oligonucleotides as a control to the total absorption. This gave insights on the ability of the aptamer to hybridize to its complementary strand and the effect of steric hinderance.

### Aptamer quantification using MiION®-nanopores

Prior to use, the flongle flow cells, were brought to room temperature on a desk for about 30 minutes. Afterwards the flow cell was inserted to the MinION® using the flongle adapter. A flow cell check was proceeded, to check the number of pores available for sequencing. The flow cell was washed with 120 µl of ONT-flush buffer and 30 µl sample, were added dropwise. Sequencing run was started using the following settings: LSQK-109 sequencing kit, no barcoding, 30 minutes runtime and bulkfile output with raw data. After use, flow cells can be re-used several times (approximately 5-times) by washing with 120 µl ONT-flush buffer between the runs. For aptamer calibration, the event-density of 0-10 µM ethanolamine aptamer in ONT-flush buffer was measured using our evaluation software. The linear range was used for aptamer quantification in the strand-displacement assay.

### Determination of signal density

The procedure to calculate the event densities was developed based on extensive data mining where data of 52 runs with known concentrations was used to first establish said procedure and then determine a calibration curve. The high data redundancy provided by the flow cells allows for robust filtering of the data prior to signal density calculation, which was also established through the same process. A combination of several pre-processing steps - including filtering and noise reduction/smoothing – and the choice of an appropriate event band was analysed with respect to their feasibility as indicator for the concentration. Hypothesizing a monotone relationship between the events and the concentration the combination of pre-processing and event band yielding the highest spearman correlation for that dependency was determined. In a first step, the data of each channel was divided in several bands, compatible to the three stages of the pore as visualized in Figure 1*A* (baseline, event, closed; closed pore not shown). This also includes determination of the baseline value – as those already fluctuate between different channels in the same run, and even more so between runs – and accompanying filtering of the channels as active or not (on average 55 of the 126 channels of a flow cell were found active, with 45 being left after filtering).

For each band the amount of datapoints in this band was determined, reflecting the share of the whole run in the according stage. Channels were then rejected from further analysis when the baseline value or baseline share of the channel are considered outliers according to a Huber-type skipped mean rule [18] (i.e. if the deviate more than 3 times the median absolute deviation from the median). The final signal density was than calculated as mean signal density over all non-rejected channels.

### Visualizing experiments with Nanotrace

Nanotrace, a python-based GUI-Application was developed to visualize the results of a run and allow for easier calculation of the associated event densities. The application is Open Source and available for download at http://github.com/simanjo/nanotrace. It consists of three main components, an overview page for a currently loaded run, a trace viewer to display squiggle plots of single channels and a density viewer showing kernel density estimates for either single channels or several channels overlaying each other to display the distribution of a complete run at once. Numpy [20] is used to provide efficient data structures and for all statistical computations done besides the Kernel density Estimations which are performed using the statsmodel [46] package and dealing with the bulkfiles is done using h5py [6], a python interface to the HDF5 data format.

## Declarations

### Ethics approval and consent to participate

Not applicable

### Consent for publication

Not applicable

### Availability of data and materials

All data generated or analysed during this study are included in this published article [and its supplementary information files].

### Competing interests

The authors declare that they have no competing interests.

### Funding

This research was funded by the Ministry of Economics, Labor and Housing Baden-Württemberg, Germany, TechPat nano, grant number 35-4223.10/17.

## Notes

### Competing Interest Statement

The authors have declared no competing interest.

