## Supplement for "Ready-to-use nanopore platform for ethanolamine quantification using an aptamer-based strand displacement assay"

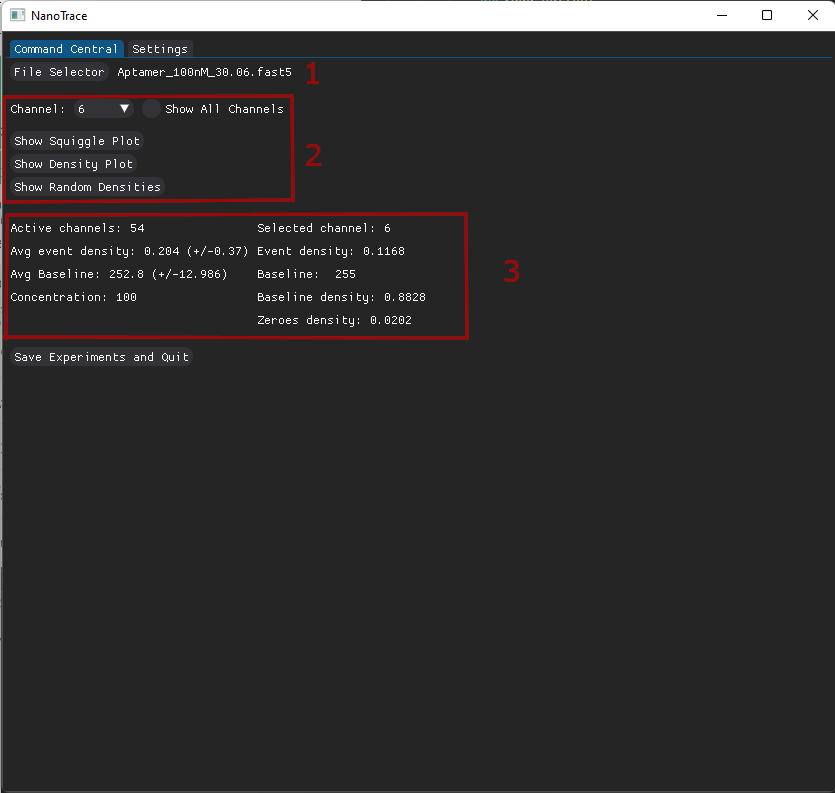

Figure 1: Main view of NanoTrace with some information for the currently loaded experiment.

Figure 1 shows the main view of NanoTrace with a fully loaded experiment. At 1, a bulkfile can be chosen to analyse/display. NanoTrace can keep a database of already loaded experiments (where files are identified via file content hash) which can be chosen in the Settings tab. If an experiment is not known, the active channels have to be calculated first (not shown). After determination of the active channels, a channel can be selected (2) and a summary is displayed (3). For this summary the name of the file is parsed to try to detect some concentration string. If the name contains “buffer” the concentration is assumed to be zero, otherwise a (case-insensitive) match for “micro”, “nano” or “nm” is performed, where prepending numbers are interpreted as concentrations in micromolar or nanomolar. Moreover, for the chosen channel a squiggle plot (Figure 2), a plot for kernel density estimates (kde) for the single channel or an overlay of 10 randomly chosen active channels kdes (Figure 3) can be displayed (2). If any of the values for zero density, baseline density or baseline value is considered an outlier in regard to this run (deviating more than 3*MAD from the median as described in the methods section) the corresponding value is printed in red.

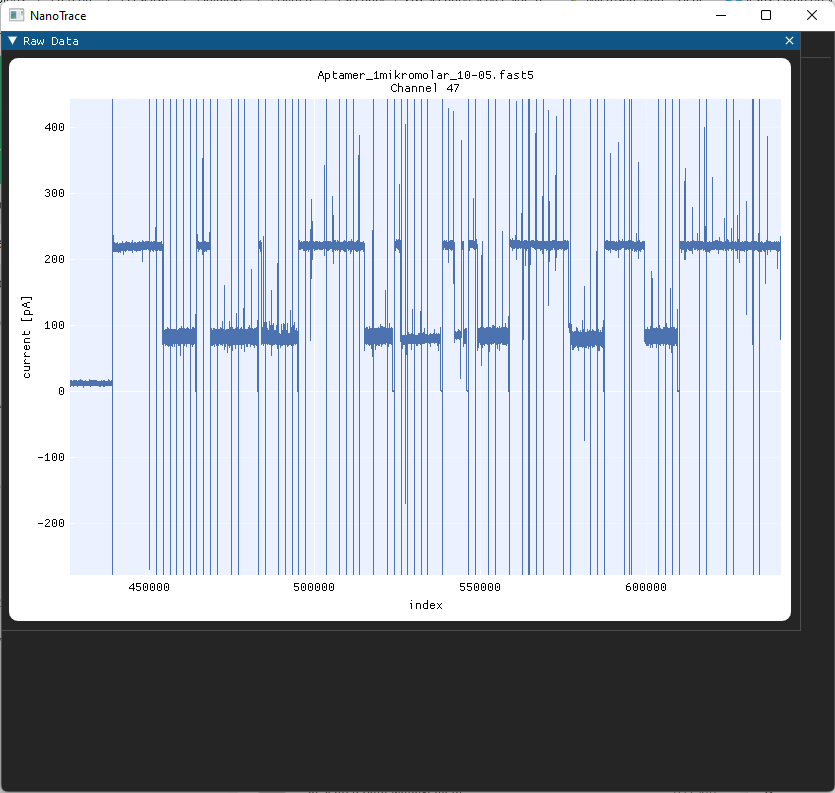

Figure 2: Squiggle plot of a single channel.

Figure 2 shows a portion of a squiggle plot for a single channel. The plot can be moved around and is freely zoomable preserving the axis ratio (scrolling inside of the plot) or each axis on its own (scrolling on the axis). The x-axis scale can be changed to show seconds (according to the sample rate recorded in the bulkfile) and the currently chosen event band can be toggled in the Settings tab.

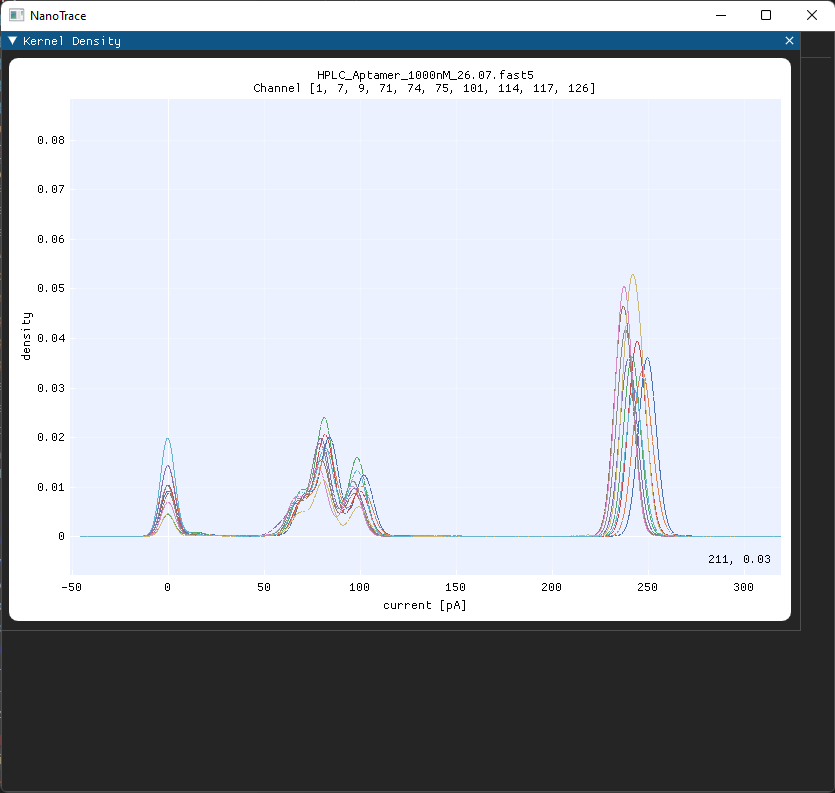

Figure 3: Kernel density estimates for 10 channels in a single plot.

Figure 3 shows 10 kdes for randomly chosen channels. The plot is similar zoomable as the squiggle plots. The amount of kdes to overlay can be changed in the Settings tab, however due to computational requirements higher values might not work too well.

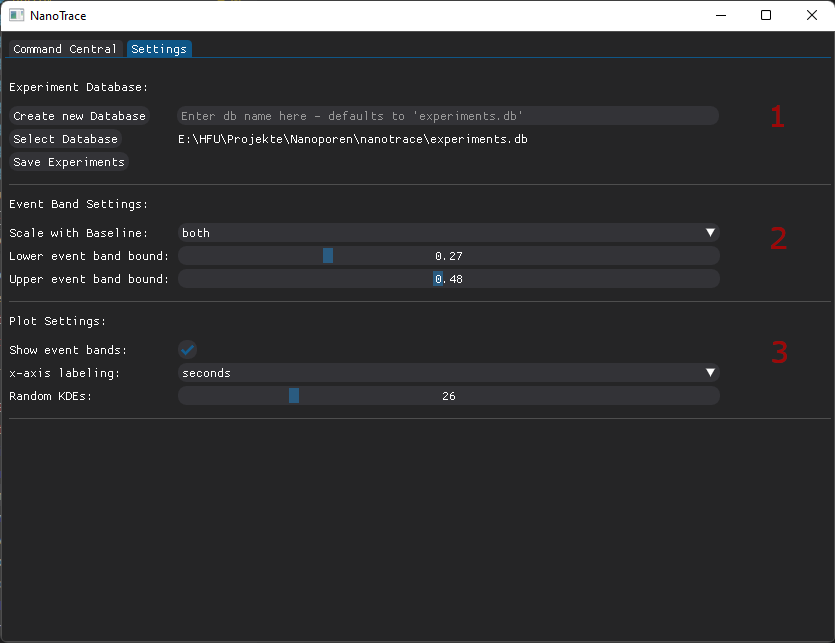

Figure 4: Settings tab to customize plotting and evaluation.

Figure 4 shows the Settings tab, used to customize the user experience.

Section 1 allows to create, select (and load) and save a file to store intermediate results for experiments. The runs are uniquely identified via their file content hash (to allow for arbitrary renaming) and the information stored are the active channels and for each the signal distribution (event band, baseline band and zero band shares) for the currently selected event band (section 2). Moreover, the current settings are also stored (and retrieved upon selection) but the band information is kept for other settings (as long as it was calculated).

Section 2 allows to customize the event band. While we found it useful to scale the event band relative to the baseline (in default parameters the event band is between 0.27*baseline and 0.48*baseline) the scaling can be adapted to only the loer or upper boundary or even completely disabled, resulting in respective absolute boundaries (in picoampere). Note however, that from theory, the drop should be scaling with the baseline, so deviating settings might not prove useful. When the band settings are changed, a recalculation of the densities has to be performed manually in the Command Central tab (to avoid lengthy background computations without explicit consent of the user).

Section 3 contains some settings for the plots. The currently selected event bands can be toggled to be shown in the squiggle plot (without a need of recalculation of the respective densities), the axis labelling of the squiggle plots can be changed between datapoints (i.e., indizes) and seconds, and the amount of random kdes to print can be set (note however that big settings might result in performance issues).
